# Elucidation of the antiviral mechanism of cystine and theanine through transcriptome analysis of mice and comparison with COVID-19 gene set data

**DOI:** 10.1101/2020.06.25.149427

**Authors:** Akira Mitsui, Ryosei Sakai, Kiyoshi Miwa, Susumu Shibahara, Shigekazu Kurihara, Kenji Nagao

## Abstract

We previously showed that oral administration of cystine and theanine (CT) to mice confers resistance to influenza virus infection. In human studies, CT prevented colds in healthy subjects and enhanced antibody production after influenza vaccination in elderly individuals with a poor nutritional status. The mechanism of action of CT is thought to be glutathione (GSH)-mediated regulation of intracellular redox, which might affect innate immune systems such as macrophages to exert physiological effects. The effect of CT on influenza is independent of viral type, and this treatment has a broad range of antiviral activities. To explore the mechanisms of CT in viral infection, we performed transcriptome profiling of spleen tissues isolated from influenza A virus (IAV)-infected mice. We identified unique gene signatures in response to CT in the IAV-infected mice. Genes upregulated by CT included redox-regulated genes such as GCLC/GCLM (subunits of glutamate cysteine ligase, a rate-limiting enzyme of GSH biosynthesis), TXN1, TXN2, TXNRD2, and SOD1, suggesting that the intracellular redox environment is substantially altered by CT. However, genes downregulated in response to CT included chemokine/chemokine receptor genes (CCL5, CCL19, CXCL9, CXCL12, CXCR3, CXCR4, and ACKR3), some of which are related to cytokine storm. A comparison with public COVID-19-related gene set data showed that the upregulated gene signature was highly similar to the downregulated gene sets of SARS-CoV/SARS-CoV-2-infected cells and the upregulated gene set of attenuated SARS-CoV-infected cells. In conclusion, the unique gene signatures observed in response to orally administered CT in IAV-infected mouse spleen tissues suggested that CT may attenuate viral infection, replication and associated symptoms such as cytokine storm.

## INTRODUCTION

Cystine, the oxidized dimer of the sulfur-containing amino acid cysteine, is a precursor of glutathione (GSH), which is responsible for the antioxidant response. Theanine is an amino acid abundant in green tea that is metabolized to glutamic acid and ethylamine. We previously showed that oral administration of cystine and theanine (CT) in mice promotes GSH synthesis and confers resistance to influenza virus infection[1]. In human studies, CT prevented colds in healthy subjects[2] and enhanced antibody production after influenza vaccination in elderly individuals with a poor nutritional status[3]. Clinical studies involving athletes showed that CT suppressed excessive inflammatory reactions induced by severe stress, such as those due to intense exercise training, and prevented a decline in immune functions[4]. The mechanism of CT has been described previously: GSH-mediated regulation of intracellular redox acts on innate immune systems such as macrophages to exert physiological effects[5][6]. The effect of CT on influenza is independent of viral type, and this treatment has been shown to increase antibody titers following hepatitis B vaccination (unpublished data), suggesting that CT has a broad range of antiviral effects.

COVID-19, caused by infection with SARS-CoV-2, is related to various physiopathological mechanisms. Numerous studies have described abnormal levels of the following cytokines and chemokines in patients (“cytokine storm”): IL-1, IL-2, IL-4, IL-6, IL-7, IL-10, IL-12, IL-13, IL-17, M-CSF, G-CSF, GM-CSF, IP-10, IFN-γ, MCP-1, MIP 1-α, HGF, TNF-α, and VEGF[7]. Among these, several cytokines are involved in Th17-type responses[8]. The key factor in SARS-CoV-2 infection could be the depletion of antiviral defenses related to the innate immune response as well as the elevated production of inflammatory cytokines[7][9]. Innate immune sensing serves as the first line of antiviral defense and is essential for immunity to viruses. The virus-host interactions involving SARS-CoV-2 likely recapitulate many of those involving other coronaviruses, given the shared sequence homology among coronaviruses and the conserved mechanisms of innate immune signaling[10].

Given its bioactivity, CT is expected to show antiviral effects against various viruses, including SARS-CoV-2. The effects of CT on the dysregulated cytokine and immune responses induced by COVID-19 are of interest. To explore the mechanisms of CT on viral infection, we performed transcriptome analysis of spleen tissues isolated from influenza virus-infected mice. Furthermore, we carried out a comparative analysis with recently published COVID-19-related gene set data[11].

## MATERIALS AND METHODS

### Animal experiment

Five-week-old female BALB/c mice were purchased from Charles River Japan (Yokohama, Japan). The mice were housed in specific pathogen-free conditions on a 24-h light-dark cycle and allowed free access to food and water. One week later, the mice were fed a control (CON) diet (68.3% α-cornstarch, 20% milk casein, 5% corn oil, 3.5% AIN-76 mineral mixture, 1% AIN-76 vitamin mixture, 2% cellulose, and 0.2% choline chloride) or a CT diet (0.5% L-cystine, 0.2% g L-theanine, 67.6% α-cornstarch, 20% milk casein, 5% corn oil, 3.5% AIN-76 mineral mixture, 1% AIN-76 vitamin mixture, 2% cellulose, and 0.2% choline chloride) for 14 days. Then, the mice were intranasally infected with a mouse-adapted strain of IAV/Aichi/2/68 (H3N2) (20 ID_50_ per mouse) using a micropipette under nembutal anesthesia. One or two days after the infection, the mice were euthanized, and then, the spleen tissues were dissected out and placed in RNAlater solution (Qiagen). All groups were in duplicate. The study was performed according to protocols approved by the Institutional Ethical Committee for Animal Research and Institutional Biosafety Committee of Ajinomoto Co., Inc.

### Gene expression profiling

Total RNA was extracted from the spleen tissues using the RNeasy Mini Kit (Qiagen), and RNA quality was checked using the Agilent 2100 Bioanalyzer (Agilent Technologies). RNA samples of each group were pooled, and the gene expression profile of the RNA samples was determined using GeneChip MG-U74Av2 arrays (Affymetrix) according to the manufacturer’s protocols. The pivot table including “Signal”, “Detection Call” and Affymetrix unique probe set identifier columns was obtained using Microarray Suite software.

### Bioinformatics analysis

All bioinformatics analyses were conducted using R (version 3.6.2). The probe sets for which at least one sample was not called “present” were initially filtered out. For identification of differentially expressed genes (DEGs) between different conditions, Weighted Average Difference (WAD) function in the TCC package (version 1.26.0) was used[12]. In the initial data exploration, principal component analysis (PCA) was performed using the prcomp function. Enrichment analysis was performed using the enrichR package (version 2.1)[13]. UpSet plot was constructed using the UpSetR package (version 1.4.0)[14]. The similarity of the gene sets was calculated with the Jaccard index[13]:

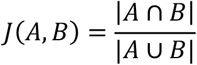

The COVID-19 Gene and Drug Set Library[11] (https://amp.pharm.mssm.edu/covid19/) was used to determine the relationships with the gene sets of SARS-CoV/SARS-CoV-2-infected cells. Among 420 gene sets (as of June, 2020), the gene sets for GSE147507[9] and GSE148729 and GSE30589[15] [NCBI’s Gene Expression Omnibus (GEO) Series IDs] (https://www.ncbi.nlm.nih.gov/geo/) were selected.

## RESULTS

### Overview of transcriptome data

We filtered out probe sets in which at least one sample was not called “present”. We selected 5,812 probes of 12,488 total probes. PCA showed three clusters, d0 (CON or CT diet, just before the infection), IAV-CON (CON diet, one or two days after IAV infection), and IAV-CT (CT diet, one or two days after IAV infection) (Fig. 1A). We then identified DEGs between these groups. DEG identification was performed using the WAD algorithm[12] with the WAD statistic (wad) set to > 0.1 or < −0.1. The comparison of IAV-CON vs. d0 yielded 978 DEGs (“IAV-CON_up”, 523 upregulated genes; “IAV-CON_down”, 455 downregulated genes). The comparison of IAV-CT vs. IAV-CON yielded 1,290 DEGs (“IAV-CT_up”, 660 upregulated genes; “IAV-CT_down”, 630 downregulated genes) (Fig. 1B, Table S1).

**Fig. 1.**
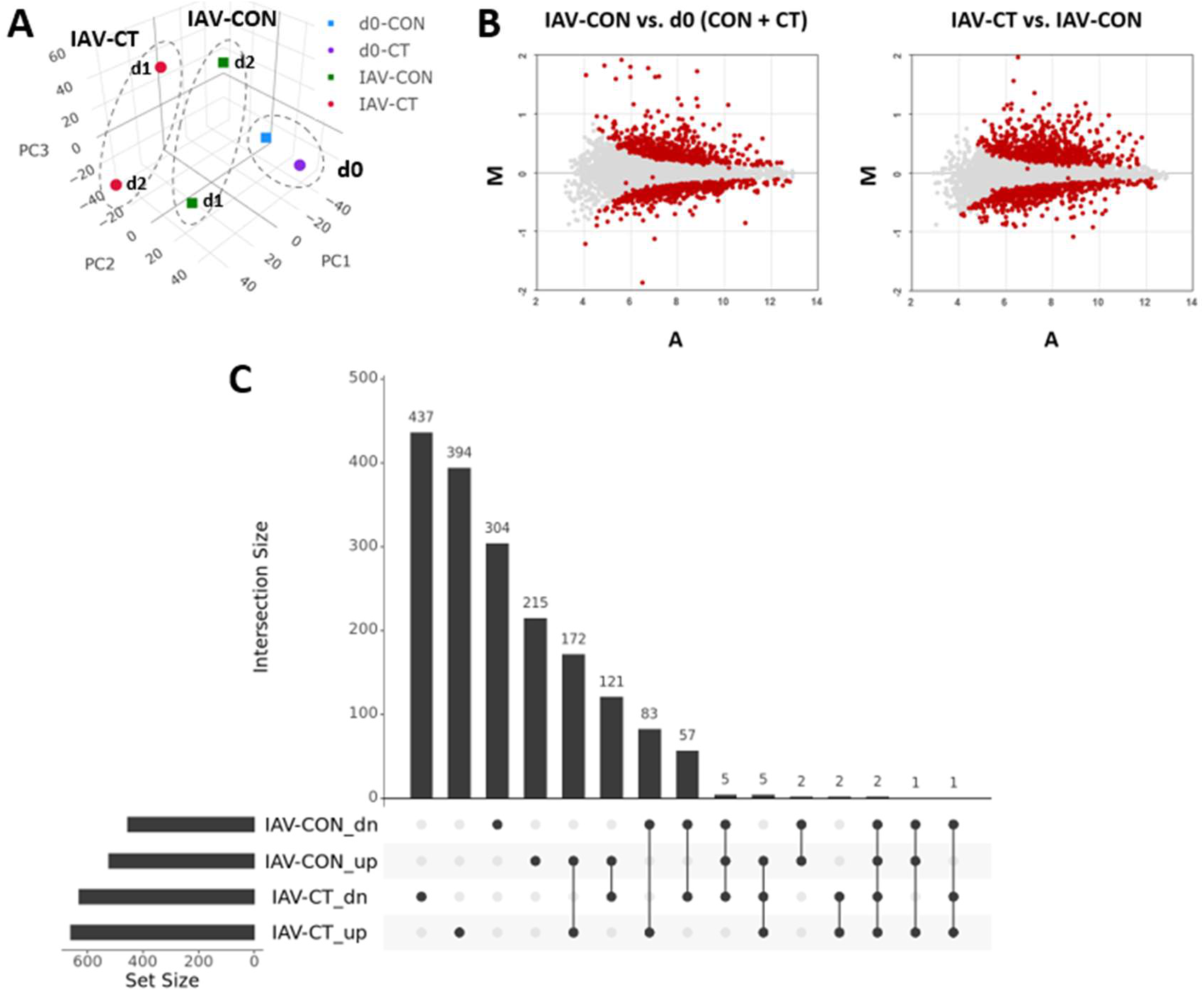
Transcriptome analysis of spleen tissues from the CT-fed mice after IAV infection. (A) PCA of the microarray data. Three clusters, d0 (CON or CT diet, just before the IAV infection), IAV-CON (CON diet, one or two days after IAV infection) and IAV-CT (CT diet, one or two days after IAV infection), are shown. (B) MA plots of the microarray data. The logged intensity ratio (M) versus the mean logged intensities (A) were plotted. DEGs (|WAD statistic| > 0.1) are indicated in red. (C) UpSet plot for the number of DEGs. IAV-CON_dn, downregulated in IAV-CON vs. d0 (CON + CT); IAV-CON_up, upregulated in IAV-CON vs. d0 (CON + CT; IAV-CT_dn, downregulated in IAV-CT vs. IAV-CON; IAV-CT_up, upregulated in IAV-CT vs. IAV-CON.

### Gene expression changes in response to CT

Among the 660 IAV-CT_up genes, 180 genes (172 + 5 + 2 + 1, Fig. 3C) were also upregulated in response to IAV infection (“IAV-CON_up”); however, 480 genes were specifically upregulated in response to CT. Both glutamate cysteine ligase catalytic subunit (GCLC) and glutamate cysteine ligase modifier (GCLM) were upregulated in response to CT (Table S1). Redox-related genes such as TXN1, TXN2, TXNRD2, PRDX2, PRDX3, PRDX5 and SOD1 were also upregulated in response to CT (Table S1).

Among the 630 IAV-CT_down genes, 65 genes (57 + 5 + 2 + 1, Fig. 3C) were also downregulated in response to IAV infection (“IAV-CON_down”); however, 565 genes were specifically downregulated in response to CT. Chemokine/chemokine-receptor genes (CCL5, CCL19, CXCL9, CXCL12, CXCR3, CXCR4, and ACKR3), MHC class I genes (H2-D1 and H2-K2), and MHC class II genes (H2-Ab1, H2-Eb1, H2-DMb1 and H2-DMb2) were downregulated in response to CT (Table S1).

### Enrichment analysis

Enrichment analysis was performed using the R package enrichR[13]. The Enrichr libraries consisted of 166 gene set libraries and 335,434 annotated gene sets (terms) as of June, 2020 (https://amp.pharm.mssm.edu/Enrichr/). When the adjusted P-value was set to less than 10^−7^, 21,567 terms were extracted: 6,055 for IAV-CON_up, 3,278 for IAV-CON_down, 9,932 for IAV-CT_up, and 2,302 for IAV-CT_down gene signatures.

When the 21,567 terms were filtered with the libraries “Gene_Perturbations_from_GEO_up” or “Gene_Perturbations_from_GEO_down”, the 10 most significant (based on the adjusted P-value) terms for each input gene sets (deg_id) were selected, as shown in Table S2. The terms for IAV-CT_up and IAV-CON_up were similar, and the intersections between the overlapping genes for IAV-CT_up and IAV-CON_up were nearly half or more than half of them (Fig. 2A). However, the enrichment for IAV-CT_up was more significant than that for IAV-CON_up (see the adjusted P-value in Table S2), and the numbers of IAV-CT_up only genes were much greater than that of the IAV-CON_up only genes (Fig. 2A), suggesting that CT enhances some signals associated with IAV infection and results in additional signals. The biological significance of genes that were upregulated by CT alone is an issue for further study. However, the intersections between the overlapping genes for IAV-CT_down and IAV-CON_down were smaller than that described above (Fig. 2B). This finding may indicate that CT downregulated unique signals that differ from those affected by IAV.

**Fig. 2.**
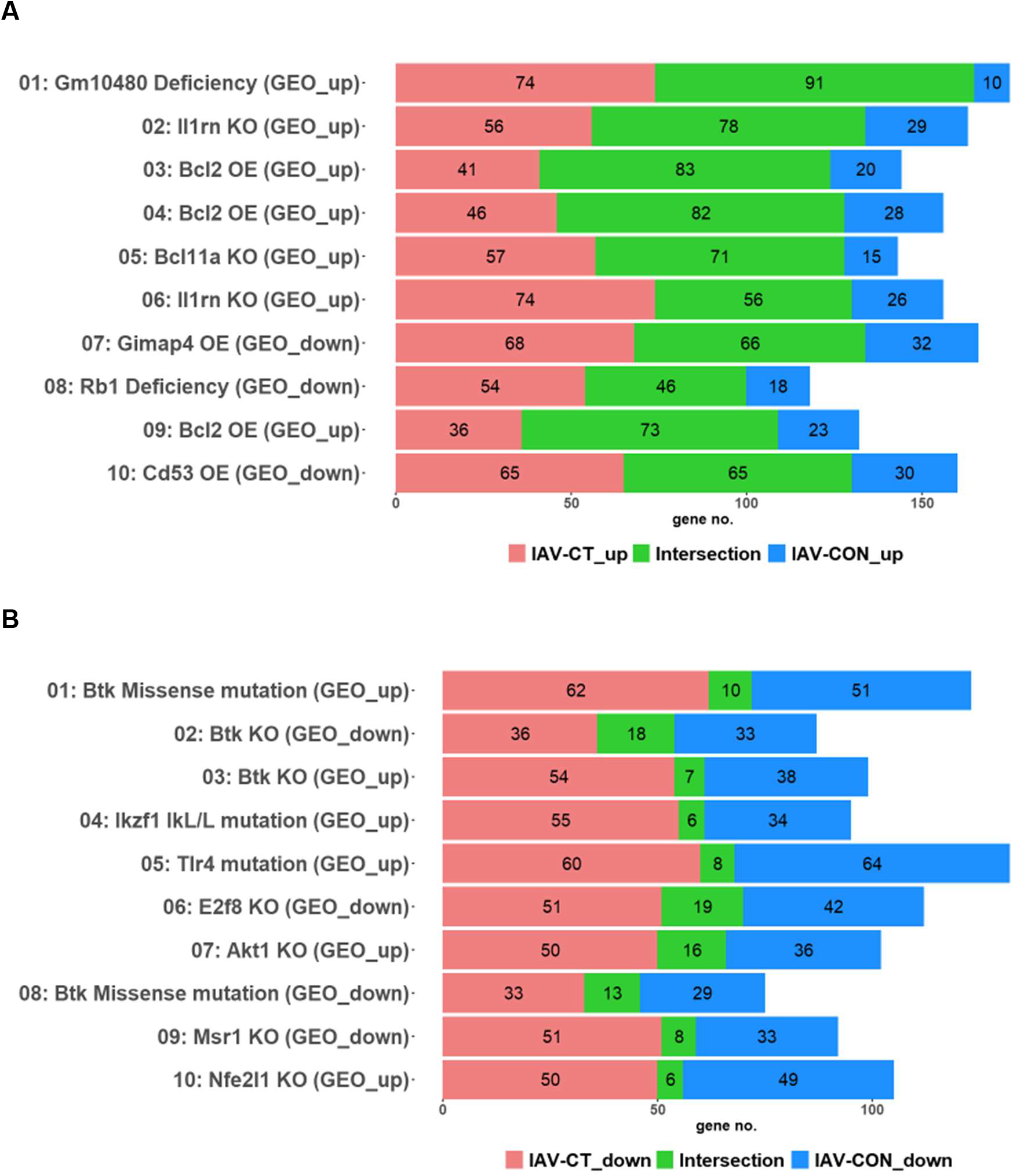
Comparison of Enrichr analysis results. (A) Top 10 Enrichr results for the IAV-CT_up signature and “Gene_Perturbations_for_GEO_up/down” libraries were compared with the IAV-CON_up results. (B) Top 10 Enrichr results for the IAV-CT_down signature and “Gene_Perturbations_for_GEO_up/down” libraries were compared with the IAV-CON_down results. Color bars indicate overlapping genes only for each signature (“IAV-CT_up” etc.) or shared genes (“Intersection”). GEO_up: Enrichr terms in “Gene_Perturbations_for_GEO_up”; GEO_down: Enrichr terms in “Gene_Perturbations_for_GEO_down”

**Fig. 3.**
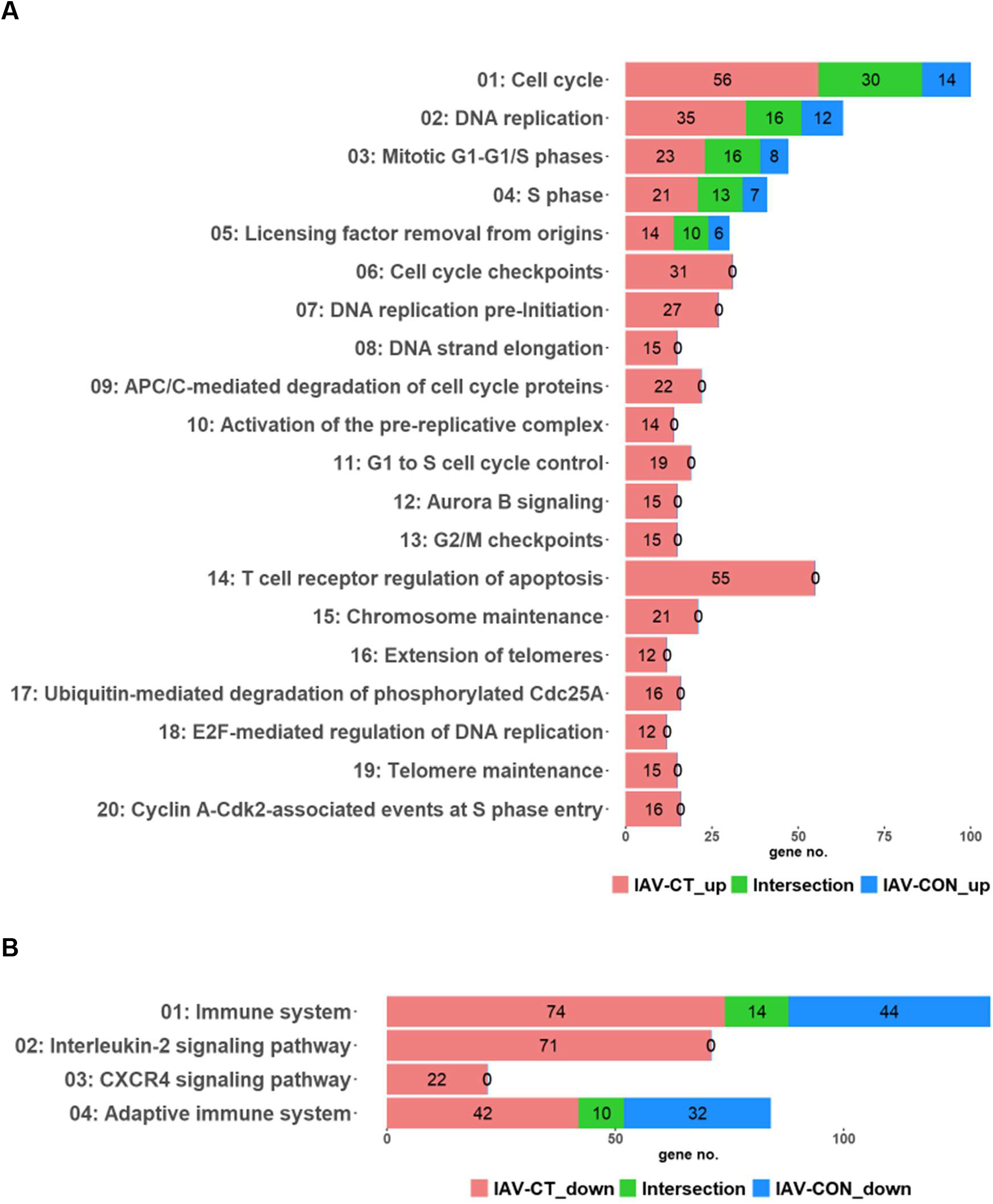
Comparison of Enrichr analysis results. (A) 20 Enrichr results for IAV-CT_up signature and “BioPlanet_2019” library were compared with the IAV-CON_up results. (B) 4 Enrichr results for IAV-CT_down signature and “BioPlanet_2019” library were compared with the IAV-CON_down results. Color bars indicate overlapping genes only for each signature (“IAV-CT_up” etc.) or shared genes (“Intersection”).

When the 21,567 terms were filtered with the library “BioPlanet_2019”, 36 terms were extracted (Table S2). The National Center for Advancing Translational Sciences (NCATS) BioPlanet is a comprehensive, publicly accessible informatics resource that catalogues all pathways, their healthy and disease state annotations, and targets within and relationships among them[16]. All 20 terms for IAV-CT_up were in the categories of “DNA replication” and/or “cell division”. Pathways containing “degradation” appeared only for IAV-CT_up (“APC/C-mediated degradation of cell cycle proteins” and “Ubiquitin-mediated degradation of phosphorylated Cdc25A”, see Fig. 3A). However, pathways related to “interferon signaling” or “immune system” were only for IAV-CON_up (“Interferon signaling”, “Immune system”, and “Immune system signaling by interferons, interleukins, prolactin, and growth hormones”) (Table S2). For the downregulated gene signatures, “Interleukin-2 signaling pathway” and “CXCR4 signaling pathway” were related to IAV-CT_down but not IAV-CON_down (Fig. 3B). Overlapping genes in these terms for the IAV-CT_down signature contained chemokine/chemokine receptor genes, such as CCL5, CXCL12, CXCR3, and CXCR4 (Table S2).

### Comparison with gene set data of SARS-CoV/SARS-CoV-2 infected cells

We used the COVID-19 Gene and Drug Set Library[11] to determine the relationships between gene signatures obtained in the current study and virus-infected gene signatures. The IAV-CT_up signature was highly similar to the downregulated gene sets of SARS-CoV and SARS-CoV-2-infected cells[9] (“01: SARS-CoV-2, Calu-3, down”, “02: SARS-CoV-2, A549, down” and “03: SARS-CoV-2, A549, down” in Fig. 4A; “01: SARS-CoV-1, Calu-3, down”, “02: SARS-CoV-2, Calu-3, down”, “03: SARS-CoV-1, Calu-3, down” and “04: SARS-CoV-2, Calu-3, down” in Fig. 4B). However, the IAV-CT_down signature was somewhat similar to upregulated gene sets of SARS-CoV-2-infected cells (Fig 4A, B). Shared genes with the three most similar gene sets (“09: SARS-CoV-2, A549, up” and “11: SARS-CoV-2, Calu-3, up” in Fig. 4A; “05: SARS-CoV-2, Calu-3, up” in Fig. 4B) included interferon stimulated genes (ISGs)[17], such as IFIT2, IFITM3, MX1 and MX2 (see “GSE147507” and “GSE148729” sheets in Table S3). Similar to the results above, the IAV-CT_up signature was highly similar to downregulated gene sets of SARS-CoV-infected Vero E6 and MA-104 cells[15] (Fig. 4C). Interestingly, the IAV-CT_up signature was also highly similar to an upregulated gene set of attenuated virus-infected cells (“04: ΔE/SARS-CoV”, rSARS-CoV-ΔE-infected compared to rSARS-CoV-infected MA-104 cells). Consistent with the characteristics of rSARS-CoV-ΔE-infected cells[15], the upregulated genes included heat shock protein (HSP) genes such as HSP90AA1, HSPD1, UBE2S, DNAJB1, and DNAJB2 (see “GSE30589” sheet in Table S3).

**Fig. 4.**
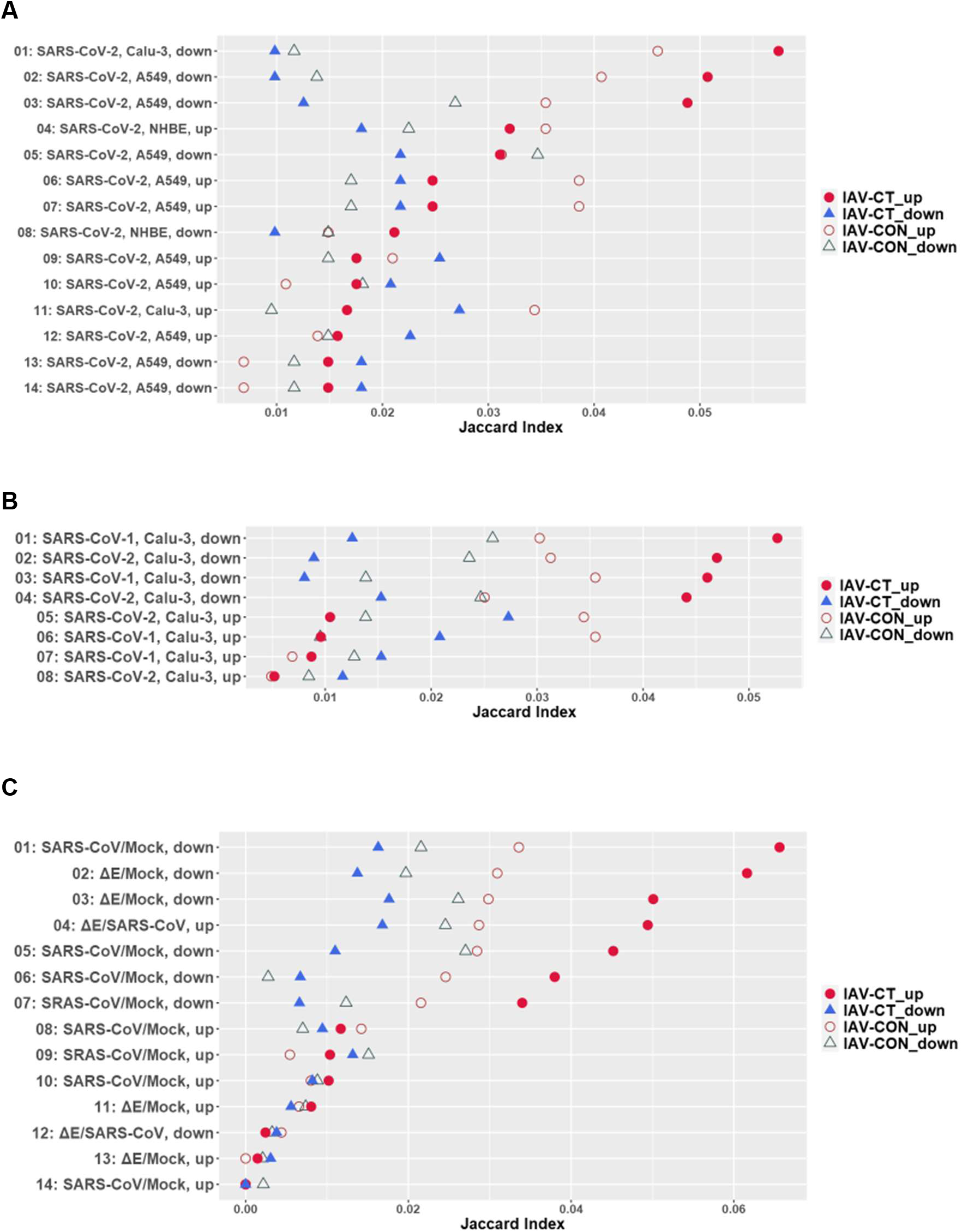
Similarity with COVID-19 gene set data. Gene set data of GSE147507 (A), GSE148729 (B) and GSE30589 (C) in the COVID-19 Gene and Drug Set Library[11] were compared with the IAV-CT_up (filled circles), the IAV-CT_down (filled triangles), the IAV-CON_up (open circles) and the IAV-CON_down (open triangles) gene signatures. Similarity between gene sets are indicated as Jaccard index.

To summarize the results, MA plot of IAV-CT vs. IAV-CON data was overlaid with the genes mentioned above (Fig. 5).

**Fig. 5.**
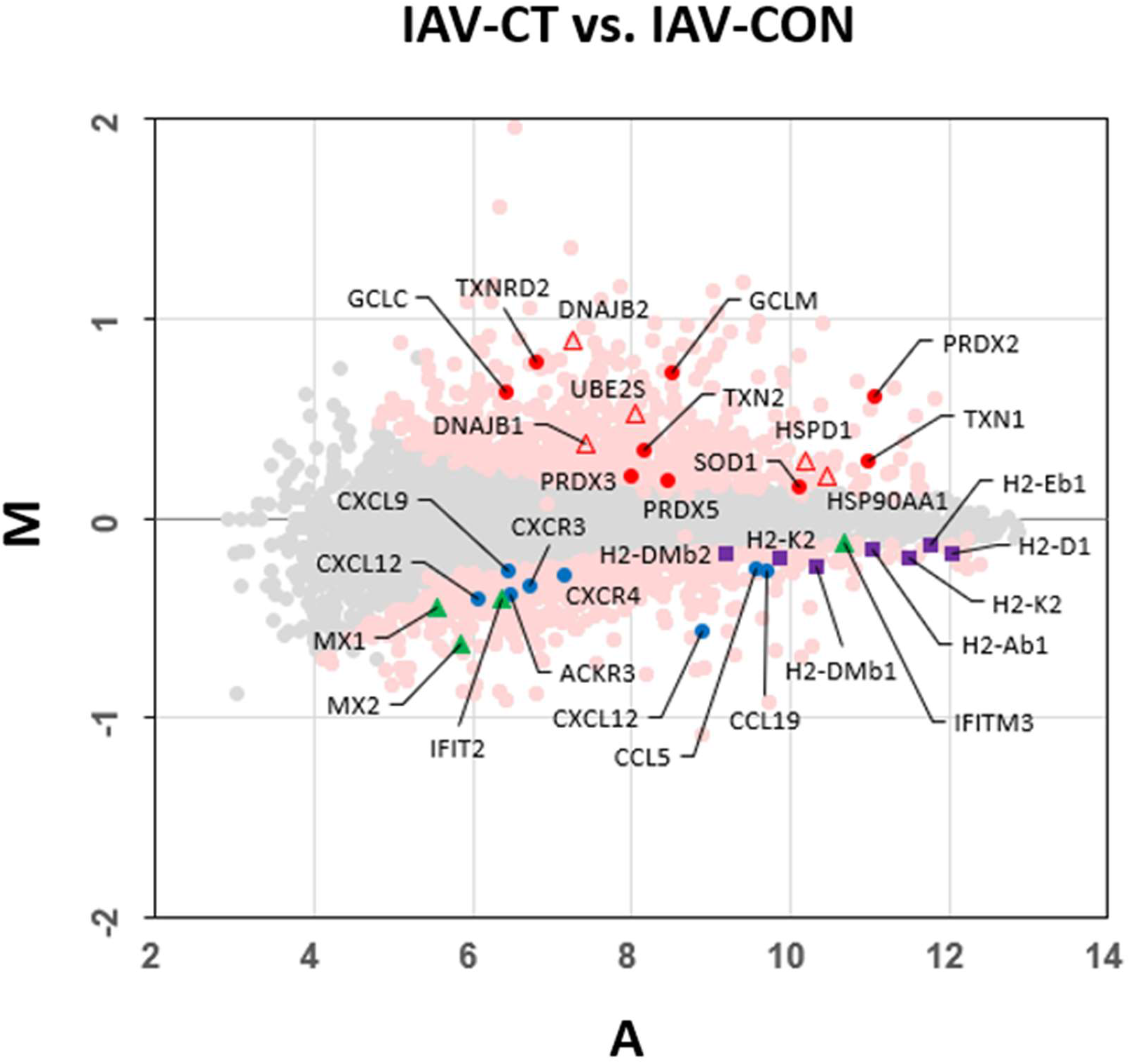
MA plot overlaid with the genes mentioned in the Results. MA plot of IAV-CT vs. IAV-CON data (see Fig. 1B) is overlaid with redox-related genes (red circles), chemokine/chemokine receptor genes (blue circles), HMC class I/II genes (purple squares), ISGs (green triangles) and HSP genes (red open triangles).

## DISCUSSION

Here, we demonstrated the molecular mechanism of the antiviral effect of orally administered CT and further explored how CT acts on the dysregulated cytokine and immune response characteristic of COVID-19. Unique gene signatures induced by CT in the IAV-infected mice, including upregulation of redox-related genes and downregulation of genes related to chemokines, were identified. A comparison with publicly available gene set data demonstrated that CT may attenuate viral infection and replication, including that of IAV, SARS-CoV, and SARS-CoV-2.

Our transcriptome analysis of spleen tissues obtained from the IAV-infected mice revealed that both GCLC and GCLM were upregulated in response to CT (Table S1). Glutamate cysteine ligase catalytic subunit (GCL), a rate-limiting enzyme of GSH biosynthesis, is composed of catalytic (GCLC) and modifier (GCLM) subunits[18]. Thus, the administration of CT not only provides substrates but also upregulates genes for GSH synthesis. Furthermore, redox-related genes such as TXN1, TXN2, TXNRD2, PRDX2, PRDX3, PRDX5 and SOD1 were upregulated in response to CT (Table S1), suggesting that the intracellular redox environment is substantially altered by CT. Diotallevi et al. demonstrated that GSH fine-tunes the innate immune response to infection and showed that GSH is important in directing changes in the gene expression profile toward the activation of host defense[6]. CT may establish intracellular GSH levels and redox environments rather than directly affecting the expression of genes involved in the innate immune response.

A comparison with the COVID-19 Gene and Drug Set Library[11] showed that the IAV-CT_up gene signature was highly similar to the downregulated gene sets of SARS-CoV- and SARS-CoV-2-infected cells. The IAV-CT_up gene signature was also highly similar to the upregulated gene signature of attenuated virus-infected cells (rSARS-CoV-ΔE-infected cells compared to rSARS-CoV-infected cells). Deletion of the E gene from SARS-CoV increases the expression of host genes related to heat shock proteins (HSPs)[15]. The presence of HSPs on the cell surface facilitates the elimination of infected cells by natural killer (NK) and T cell subsets[19]. In fact, IAV-CT_up genes included HSPs such as HSP90AA1, HSPD1, UBE2S, DNAJB1, and DNAJB2, suggesting that CT might eliminate virus-infected cells by a mechanism similar to that of SARS-CoV-ΔE. E protein is also common in other respiratory viruses, including SARS-CoV-2[20]; therefore, CT may reduce coronavirus infection and replication. However, the IAV-CT_down signature was similar to the upregulated gene sets of SARS-CoV-2-infected cells, and the overlapping genes included ISGs (IFIT2, IFITM3, MX1 and MX2). Since the elevated expression of ISGs is thought to be the result of viral infection/replication, the response observed in IAV-CT_down also suggests that CT could attenuate viral infection and/or replication.

In this study, genes encoding chemokines, some of which are characteristic of cytokine storm, were downregulated in response to CT. Among these molecules, CCL5 and CXCL9 have been shown to be elevated in response to SARS-CoV-2, SARS-CoV or MERS-CoV infections[9][21]. The IAV-CT_down signature also contained CXCL12 and CXCR4, which are thought to be involved in the activation of Th17 cells[22][23].

Our study has some limitations. For instance, we compared IAV and IAV-CT samples and therefore did not observe transcriptome changes in response to CT alone. Although we showed the downregulation of chemokine genes and virus-responsive genes, these changes could be due to the reduction of IAV infection and/or replication. Further work is needed to clarify the detailed mechanisms of the antiviral effects of CT.

In conclusion, the present study demonstrated the unique gene signatures in response to orally administered CT in IAV-infected mouse spleen tissues. Upregulated signatures included redox-related genes, and downregulated genes included chemokines. Based on a comparison with COVID-19-related public gene set data, CT may attenuate viral infection, replication and the cytokine storm related to COVID-19. Future clinical work is needed to confirm these findings.

